# Monitoring the spread of SARS-CoV-2 variants in Moscow and the Moscow region using targeted high-throughput sequencing

**DOI:** 10.1101/2021.07.15.452488

**Authors:** N.I. Borisova, I.A. Kotov, A.A. Kolesnikov, V.V. Kaptelova, A.S. Speranskaya, L.Yu. Kondrasheva, E.V. Tivanova, K.F. Khafizov, V. G. Akimkin

## Abstract

Since the outbreak of the COVID-19 pandemic caused by the SARS-CoV-2 coronavirus, the international community has been concerned about the emergence of mutations that alter the biological properties of the pathogen, for example, increasing its infectivity or virulence. In particular, since the end of 2020, several variants of concern have been identified around the world, including variants “alpha” (B.1.1.7, “British”), “beta” (B.1.351, “South African”), “gamma” (P.1, “Brazilian”) and “delta” (B.1.617.2, “Indian”). However, the existing mechanism for searching for important mutations and identifying strains may not be effective enough, since only a relatively small fraction of all identified pathogen samples can be examined for genetic changes by whole genome sequencing due to its high cost. In this study, we used the method of targeted high-throughput sequencing of the most significant regions of the gene encoding the S-glycoprotein of the SARS-CoV-2 virus, for which a primer panel was developed. Using this technique, we examined 579 random samples obtained from patients in Moscow and the Moscow region with coronavirus infection from February to June 2021. The study demonstrated the dynamics of the representation in the Moscow region of a number of SARS-CoV-2 strains and its most significant individual mutations in the period from February to June 2021. It was found that the strain B.1.617.2 began to spread rapidly in Moscow and the Moscow region in May, and in June it became dominant, partially displacing other varieties of the virus. The results obtained make it possible to accurately determine the belonging of the samples to the abovementioned and some other strains. The approach can be used to standardize the procedure for searching for new and existing epidemiologically significant mutations in certain regions of the SARS-CoV-2 genome, which allows studying a large number of samples in a short time and to get a more detailed picture of the epidemiological situation in the region.

## Introduction

Since its first detection in December 2019 in Wuhan (Zhou et al. 2020), the SARS-CoV-2 coronavirus has spread worldwide and has caused nearly 4 million deaths (https://coronavirus.jhu.edu/). Starting from the beginning of the pandemic, a number of therapeutic and preventive measures to combat COVID-19 have been developed, which include the use of immunological drugs, for example, monoclonal antibodies (Chen et al. 2021; Weinreich et al. 2021) and vaccines (Baden et al. 2021; Polack et al. 2020; Jones and Roy 2021; Ryzhikov et al. 2021), for which the S-protein SARS-CoV-2 usually acts as an antigen.

At the end of 2020, the international scientific community described several SARS-CoV-2 variants of concern that require special attention. These now include “alpha” (formerly called “British”, B.1.1.7), “beta” (“South African”, B.1.351), “gamma” (“Brazilian”, P.1), and “delta” (“Indian”, B.1.617.2). These variants of the virus interested researchers after reports of an increase in cases of their transmission from person to person began to appear in a number of geographic regions, and subsequently new variants of the pathogen were discovered in many countries. For example, the alpha variant quickly spread to the Southeast of the UK, where it caused a large number of COVID-19 cases, and was identified in the US shortly thereafter (CDC, https://www.cdc.gov/coronavirus/2019-ncov/Transmission/variant.html), becoming dominant in the country by April 2021. Similarly, variants from South Africa and Brazil have triggered disease outbreaks in their respective countries. They are also of concern because they contain the E484K mutation in the spike (S-) protein, which is likely to reduce the effectiveness of some therapeutic monoclonal antibodies, making it difficult to neutralize the virus *in vitro*, and which could lead to a potential escape from immunity caused by previous infection or vaccination (Wang et al. 2021; Garcia-Beltran et al. 2021; Liu et al. 2021; Yuan et al. 2021; Ikegame et al. 2021; Ryzhikov et al. 2021). In addition, three variants (“alpha”, “beta” and “gamma”) have the N501Y mutation in the S-protein gene, which is associated with increased affinity for the angiotensin-converting enzyme 2 (ACE2) receptor, which possibly contributes to the increased transmissibility of the virus (Nelson et al. 2021; Tian et al. 2021). Earlier, specialists from the Central Research Institute of Epidemiology of Rospotrebnadzor developed a set of reagents for the rapid detection of the N501Y mutation in the virus genome using the isothermal loop amplification method (Khafizov et al. 2021), which dramatically reduced the number of samples transferred for whole genome sequencing in order to identify and monitor new strains containing the above mutation. However, the emergence of new strains, including those characterized by other mutations in the S-protein gene, showed that genomic substitutions at the sites of LAMP primer annealing can reduce the effectiveness of the technique. In addition, as SARS-CoV-2 variants are discovered, the list of mutations to be tracked is growing. For example, in Russia, the “delta” strain was discovered in May, which previously caused a high incidence in India, and its prevalence in the country has grown rapidly since then. Moreover, in Russia, local strains have also been identified, including “Siberian” (B.1.1.397 +) and “North-Western” (B.1.1.370.1), which have mutations in the S-protein gene (Gladkikh et al 2021; Klink et al. 2021). At the moment, studies of strains circulating in Russia are constantly ongoing (Komissarov et al. 2021). For these reasons, researchers need more efficient and versatile tools to identify a range of important mutations in a single analysis. Although whole genome sequencing is by far the most detailed method for genetic analysis of a pathogen (Long et al. 2021), this approach is not always the best from a financial point of view. The application of this technique is difficult in conditions of a constant increase in the number of infected people, including cases of reinfection; moreover, it may turn out to be irrational for quick epidemiological surveillance, given that the most significant changes occur in a small part of the pathogen’s genome.

In this article, we describe the identification of isolates belonging to epidemiologically important variants of SARS-CoV-2, including strains B.1.1.7, B.1.351 and B.1.617.2, using targeted high-throughput sequencing. For this, we have developed a primer panel that allows fast and efficient targeted amplification of genome fragments, while avoiding the stage of adapter ligation when preparing samples for sequencing due to the modification of oligonucleotides at the stage of synthesis. The amplified regions of the genome include the most significant from an epidemiological point of view mutations corresponding to amino acid substitutions K417T, L452R, T478K, E484K, S494P, N501Y, A570D, P681H and others, as well as deletions HV69-70 and Y144 (figure 1). This makes it possible not only to drastically reduce the cost sample preparation, but also to reduce the amount of generated data by orders of magnitude, allowing studying a larger number of virus samples. The latter is especially important when it is urgently required to establish the representation of various strains in the area under consideration. Using a new primer panel, we examined 579 random samples of the virus, selected in February-June 2021 in Moscow and the Moscow region, and showed the change in the frequencies of individual mutations and strains. The presented work further illustrates the need to intensify population, genomic and epidemiological studies to identify, track the spread and monitor new variants of viral pathogens.

**Figure 1.**
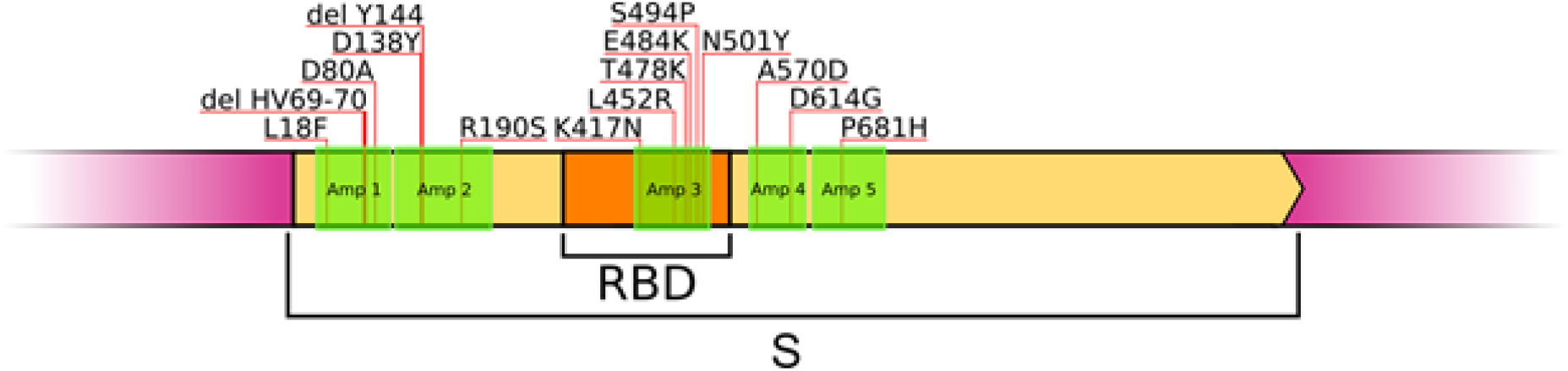
Localization of the amplicons obtained using the primer panel relative to the SARS-CoV-2 genome. Several amino acid substitutions covered by the panel are indicated.

## Methods

For the study, we used nasopharyngeal swabs from patients with symptoms of COVID-19, for whom the presence of SARS-CoV-2 was confirmed by real-time PCR with reverse transcription using the AmpliSens Cov-Bat-FL reagent kit (AmpliSens, Russia). The study was conducted with informed consent of the patients. The samples were placed in a transport medium produced by the Central Research Institute of Epidemiology. Isolation of RNA from clinical material was performed using the RIBO-prep kit (AmpliSens, Russia), reverse transcription was performed using the REVERTA-L kit (AmpliSens, Russia). Only those clinical samples were selected in which the cycle threshold (Ct) for PCR did not exceed 20.

Amplification was carried out using PCR mixture-2 blue (AmpliSens, Russia) containing Taq-polymerase on a T100 Thermal Cycler (Bio-Rad, USA). Next, the PCR product was purified from the reaction mixture using AMPureXP beads (Beckman Coulter, USA) Amplification temperature profile: (1) Denaturation at 95° C for 30 seconds; (2) 38 amplification cycles: 95° C - 30 sec., 60° C - 20 sec., 72 ° C - 60 sec.; (3) Final elongation at 72°C for 3 minutes. Indexing was performed using PCR-mixture-2 blue (AmpliSens, Russia) and EvaGreen (Biotium, USA) as a dye, on a QuantStudio 5 Real-Time PCR System (Thermo Fisher Scientific, USA). For this, index primers compatible with the Nextera XT Index Kit v2 were used (namely, N7xx - 26 possible variants and S5xx - 18 variants, for more details see Illumina Adapter Sequences, Document # 1000000002694 v16, April 2021). This double indexing improves the accuracy of sample identification and allows one to test more samples at a time. Indexing temperature profile: (1) 98°С - 30 sec.; (2) 15 cycles: 98°С - 10 sec., 65°С - 1 min. 15 sec. Then the reaction mixture was purified again. The DNA concentration was measured on a Qubit 4.0 fluorometer (Thermo Fisher Scientific, USA). Samples were dropped into a pool of equivalent concentration and the pool quality was analyzed using an Agilent 2100 Bioanalyzer system. Denaturation and calculation of the loading concentration were carried out according to the manufacturer’s instructions. Sequencing was performed on the Illumina MiSeq platform using the MiSeq Reagent Kit v2 (PE 150 + 150 or PE 250 + 250 cycles) or the MiSeq Reagent Kit v3 (PE 300 + 300 cycles), as well as on the Illumina HiSeq using the v2 Rapid SBS kit ( PE 250 + 250 cycles). One sample accounted for about 50,000 reads, which is equivalent to ~0.3% of the information obtained when using the MiSeq Reagent Kit v2 (PE 150 + 150) for sequencing or ~0.1% of the MiSeq Reagent Kit v3 (PE 300 + 300 cycles). The chosen approach allows studying a significant number of samples without excessive financial costs, while the average coverage for most samples is from 2,000 and above.

A panel of primers for targeted amplification of S-protein gene fragments was selected manually, taking into account the available information on known epidemiologically significant mutations, as well as data on conserved regions of the genome. Melting points of oligonucleotides and determination of the degrees of interactions between them were determined using the Multiple Primer Analyzer instrument (Thermo Fisher Scientific, USA). Using the blastn program (Altschul et al. 1990) the specificity of each obtained sequence was assessed for all known organisms, primarily *Homo sapiens*, the genetic material of which is present in the sample in the greatest amount, excluding many nonspecific interactions between the primer and regions of human and other organisms DNA. A total of 5 pairs of oligonucleotides were created, also containing additional adapter sequences necessary to reduce the time and cost of sample preparation. The primer structures are shown in Table 1. The lengths of the amplicons were selected so as to provide complete coverage of the target regions during their high-throughput sequencing on Illumina MiSeq platforms using the MiSeq reagent kits v2 (300 cycles), v2 (500 cycles) and v3 (600 cycles), and for Illumina HiSeq using the v2 Rapid SBS Kit (500 cycles).

**Table 1.**
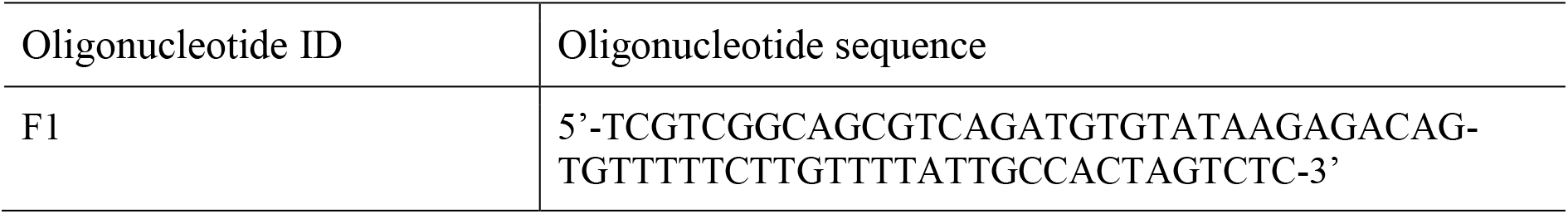
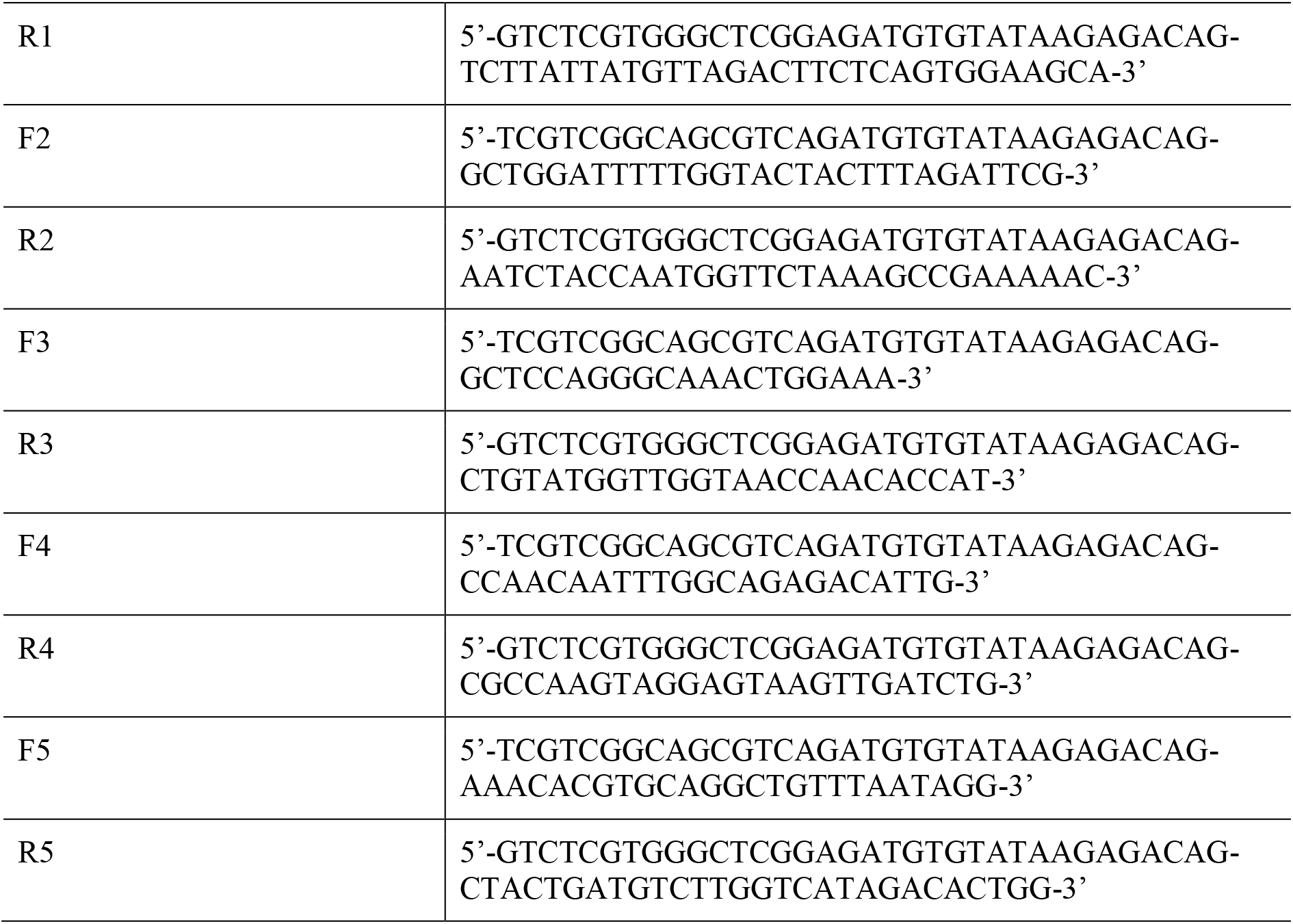
Sequences of oligonucleotides in the primer panel. The specific part is separated from the adapters with a “-” symbol.

To analyze the sequencing data, the resulting reads were aligned to the reference coronavirus genome with the bwa program (Li and Durbin 2009). Bbtools (Bushnell, Rood, and Singer 2017) was used to trim adapter sequences in reads. The search for genetic variants was carried out using the GATK package (Poplin et al. 2017). The sequences obtained in this study were uploaded to the VGARus database (https://genome.crie.ru/).

VMD software was used to visualize the S-protein molecule and create figures (Humphrey, Dalke, and Schulten 1996). A structural model of S-protein (PDB ID: 7CAB) obtained by cryo-electron microscopy (Lv et al. 2020) was used.

## Results

Using the primer panel described in the previous section, we sequenced 579 samples collected from patients with coronavirus infection from February to June 2021 in Moscow and the Moscow region. The time period during which the studies were carried out was a period when SARS-CoV-2 strains began to actively spread in Russia and the rest of the world, causing concern, which could become one of the reasons for new waves of morbidity in a number of countries.

In isolates obtained in February only a very small (~2%) proportion of the “alpha” strain was found; in March its frequency increased to ~20%, which is consistent with the data that this strain has increased contagiousness (Davies et al. 2021). However, it did not receive further widespread adoption, and its share gradually decreased, and in mid-June it fell to almost zero. Probably, this was caused by the appearance in Russia of the “delta” strain in May, which earlier, possibly, caused the increased rates of morbidity and mortality among the population of India. By mid-June, the proportion of this strain also rose sharply to 70% in Moscow, and according to data not included in the presented study, it continues to increase up to more than 90%. In addition, at the end of June, there were cases of the “delta-plus” strain, which has an additional replacement K417N, which was previously found in the “beta” strain, and also located in one of the regions of the SARS-CoV-2 genome amplified using the developed primer panel.

It is noteworthy that the “beta” variant did not receive significant distribution in Moscow, although in April its share quickly rose to ~13%, causing some concern, since there is evidence that this strain partially escapes the neutralizing effect of antibodies caused by both previous coronavirus infection and vaccination.

In addition, it is worth noting strain B.1.1.523, the proportion of which increased significantly by April. The presence of the E484K mutation indicated that, like the beta strain, it may be more resistant to the action of antibodies. In addition, this variant has changes in the spike protein: S494P substitution and a deletion of three amino acids. However, its share also fell sharply in June, when the above-mentioned “delta” variant became prevalent. At the same time, we did not find cases of infection with the “gamma” (“Brazilian”) strain.

Finally, Figure 3 shows the dynamics of the representation of variants of the virus with mutations N501Y or E484K, but not assigned to any of the above strains due to the absence of other necessary changes in the genome.

**Figure 2.**
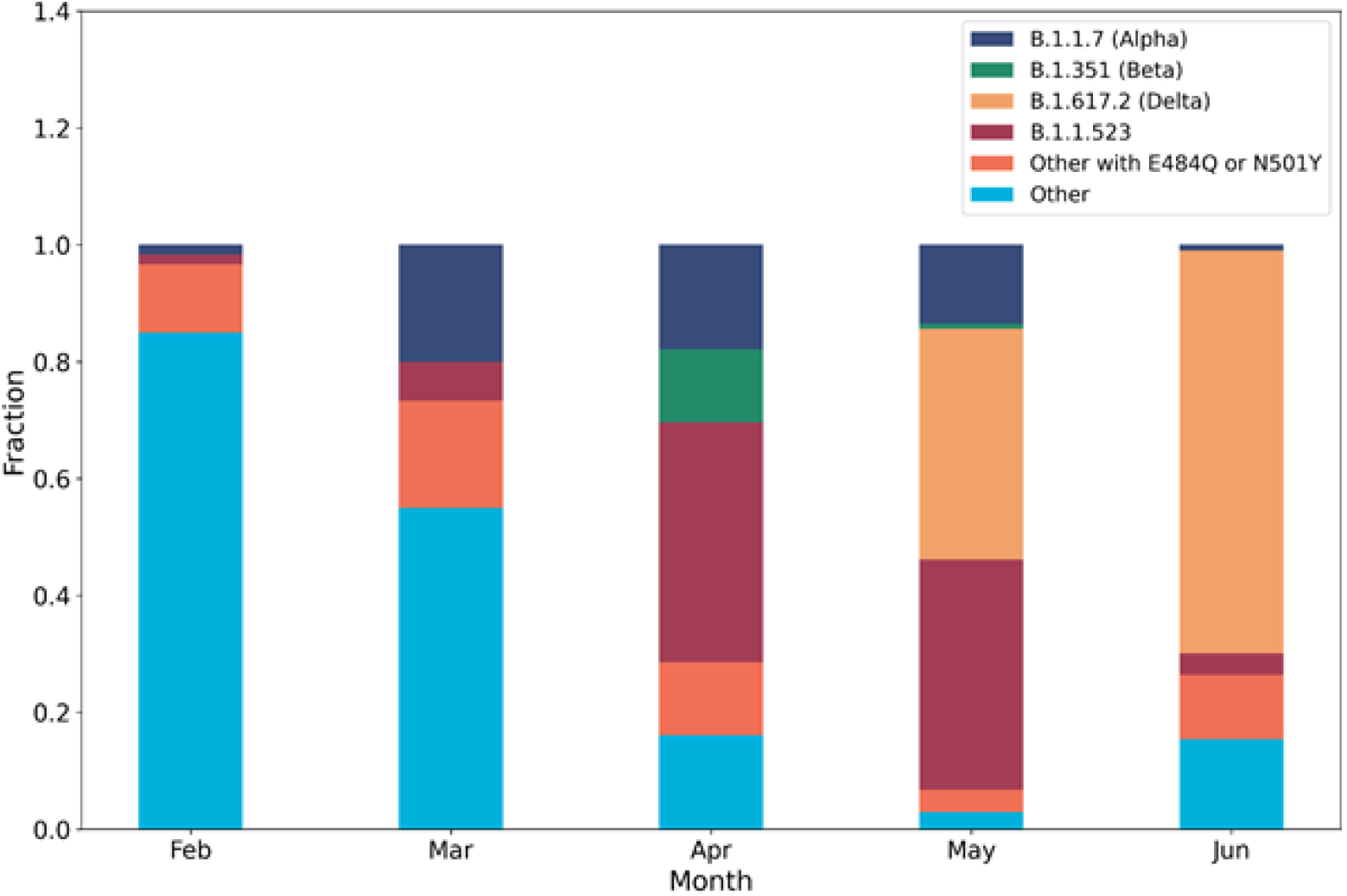
Representation of various variants of SARS-CoV-2 from February to June 2021 in Moscow and the Moscow region.

**Figure 3.**
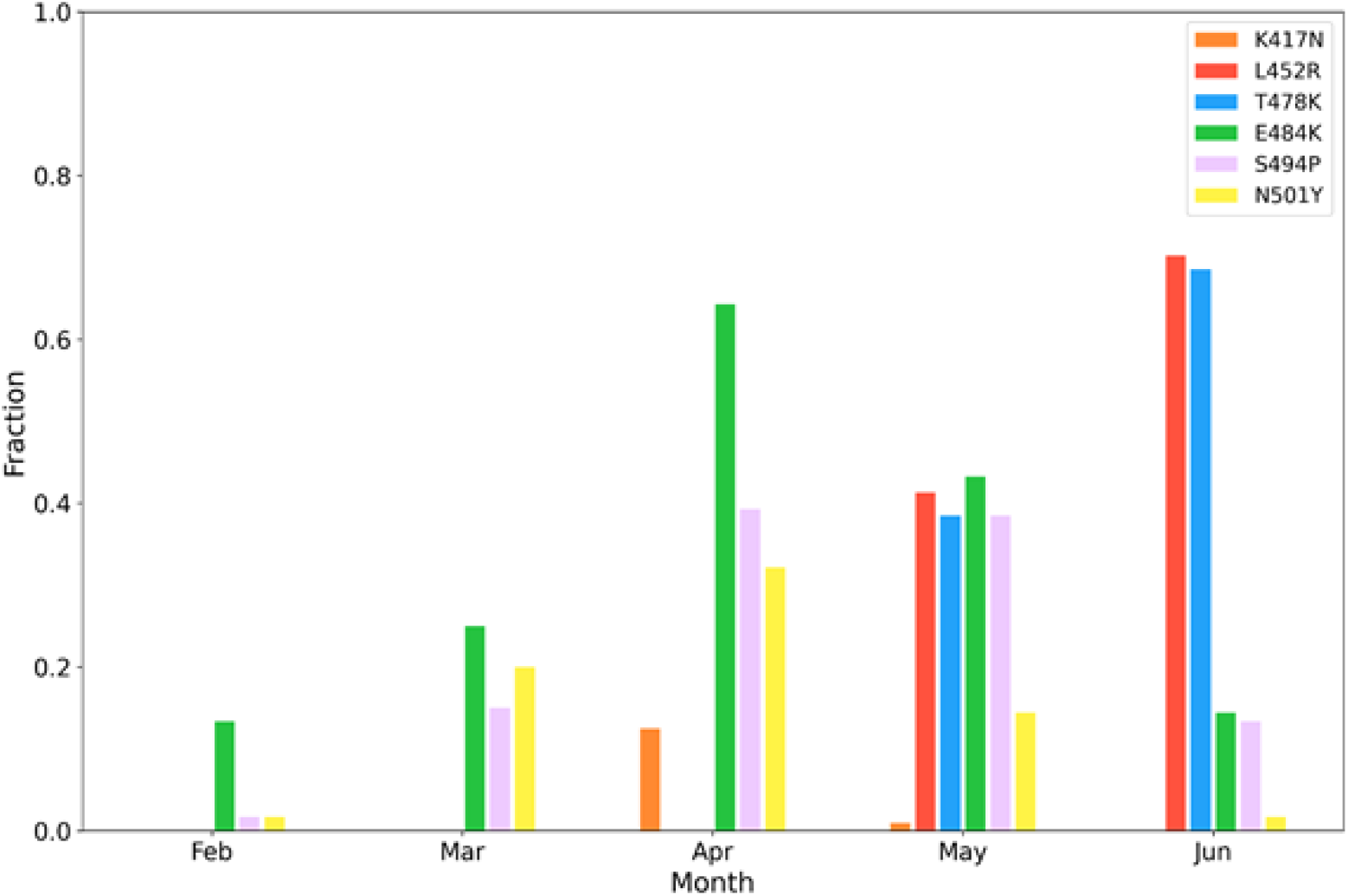
Frequencies of individual SARS-CoV-2 mutations in different months of 2021 in Moscow and the Moscow region.

The diagrams in Figure 3 demonstrate the change in the prevalence rates of individual mutations, which were previously shown to alter the properties of the virus, and lead to increased transmissibility or avoidance of the protective effect of antibodies. In February almost 15% of all isolates contained the E484K mutation, although at that time the “beta” strain containing this substitution had not yet been detected in the Russian Federation. This result shows that although such a change in the genome may give certain advantages to the virus, without other important mutations, the significant combinations of which are often not clear, does not always lead to its widespread distribution. The following months were marked by an increase in the frequencies of several mutations, incl. E484K, N501Y, S494P. In April, due to the appearance of the beta strain in the country, an increase in the frequency of the K417N mutation was noted, which is also present in the delta-plus variant that was first discovered in Russia only at the end of June. Finally, the L452R and T478K mutations belonging to the delta strain appeared in May and are now present in the overwhelming majority of the pathogen’s genomes.

The presented data show that the use of a small primer panel for amplification of fragments of the SARS-CoV-2 genome and subsequent targeted sequencing makes it possible to detect almost all known significant changes in the coronavirus S-protein gene, and to identify strains of SARS-CoV-2, making it possible to track their frequencies at regular intervals. At the same time, it should be noted that individual mutations do not always lead to a more “dangerous” variant of the virus, and only their complex combinations can give new properties to the pathogen, sometimes leading to its widespread distribution.

## Conclusion

In this methodological article, we described the possibility of detecting a number of SARS-CoV-2 strains, including variants “alpha” (“British”, B.1.1.7), “beta” (“South African”, B.1.351), “gamma” (“Brazilian”, P.1), “delta” (“Indian”, B.1.617.2), using targeted high-throughput sequencing. For this purpose, we have developed a primer panel (Table 1), which makes it possible to efficiently carry out targeted amplification of fragments of the coronavirus genome. The amplified regions of the genome include a large number of known important mutations that alter the properties of the virus, which dramatically reduces the cost of sequencing and allows studying more samples per unit of time. The latter is justified if the goal is to identify significant changes in the S-protein gene to determine the belonging of virus isolates to different strains. To demonstrate the use of the primer panel in practice, we also conducted a study of 579 random samples of SARS-CoV-2 collected in February-June 2021 from patients in Moscow and the Moscow region. It is noteworthy that most of the significant changes in the S-protein gene are localized in a number of small fragments of the gene, mainly encoding amino acids located on the surface of the protein (Figure 4).

**Figure 4.**
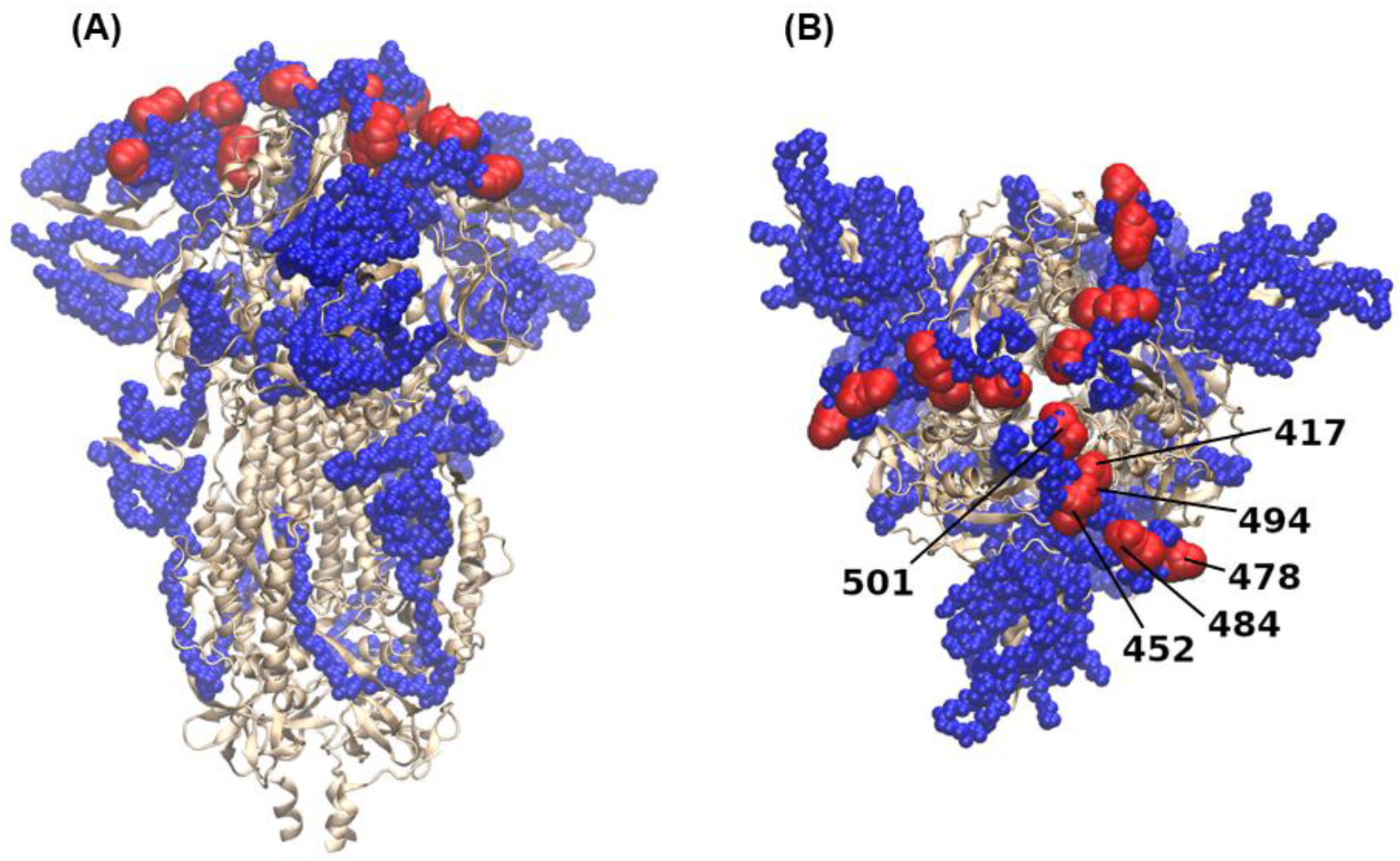
Structural model of S-protein obtained using cryo-electron microscopy (PDB ID: 7CAB; Lv et al. 2020). View (A) from the side and (B) from above. The regions included in the primer panel are highlighted in blue and shown as small spheres. Significant mutations (K417T, L452R, T478K, E484K, S494P, N501Y) are shown in red as larger spheres and are indicated in Figure (B).

We have shown a rapid change in the proportion of various genetic variants of the virus in the above period, including the emergence and active increase in the frequency of the delta strain in May-June 2021 in Moscow and the Moscow region, which may be partially responsible for the new wave of the disease in the summer. However, the population’s neglect of social distancing, personal protection and low vaccination rates also play a critical role in the spread of infection. Thus, our work further illustrates the need to intensify population, genomic and epidemiological studies to identify, track the spread and monitor new variants of SARS-CoV-2 and other viral pathogens that are characterized by increased variability.

Although we cannot exclude the possibility that new significant genomic variants will affect other important fragments of the genetic material of the virus, a panel with a small number of primers helps to dramatically reduce the cost of identifying currently circulating strains, and a simple design allows one to quickly change the structure of oligonucleotides, taking into account the constantly emerging new information about genomes. It should be noted separately that targeted sequencing cannot completely replace whole-genome sequencing, which makes it possible to identify all genomic changes and conduct a detailed phylogenetic analysis. For a detailed bioinformatics analysis of strains circulating in Russia, specialists can refer to the recently created VGARus coronavirus genome database (https://genome.crie.ru/), which contains about 15,000 sequences at the end of June 2021, including ~8,000 complete genomes obtained by sequencing isolates collected in different regions of the Russian Federation.

## Notes

**Conflict of interest**. The authors declare no conflict of interest.

### Competing Interest Statement

The authors have declared no competing interest.

